# Acoustically Detonated Biomolecules for Genetically Encodable Inertial Cavitation

**DOI:** 10.1101/620567

**Authors:** Avinoam Bar-Zion, Atousa Nourmahnad, David R. Mittelstein, Sangjin Yoo, Dina Malounda, Mohamad Abedi, Audrey Lee-Gosselin, David Maresca, Mikhail G. Shapiro

## Abstract

Recent advances in molecular engineering and synthetic biology have made it possible for biomolecular and cell-based therapies to provide highly specific disease treatment. However, both the ability to spatially target the action of such therapies, and their range of effects on the target tissue remain limited. Here we show that biomolecules and cells can be engineered to deliver potent mechanical effects at specific locations inside the body under the direction of focused ultrasound. This capability is based on gas vesicles, a unique class of air-filled protein nanostructures derived from buoyant photosynthetic microbes. We show that low-frequency ultrasound can convert these nanoscale biomolecules into micron-scale cavitating bubbles, as demonstrated with acoustic measurements and ultrafast optical microscopy. This allows gas vesicles targeted to cell-surface receptors to serve as remotely detonated cell-killing agents. In addition, it allows cells genetically engineered to express gas vesicles to be triggered with ultrasound to lyse and release therapeutic payloads. We demonstrate these capabilities *in vitro*, *in cellulo*, and *in vivo*. This technology equips biomolecular and cellular therapeutics with unique capabilities for spatiotemporal control and mechanical action.

## INTRODUCTION

Targeted molecular and cellular therapeutics have revolutionized the treatment of diseases such as cancer by allowing therapy to be aimed more specifically at pathological cells and tissues. For example, antibodies have been used to deliver molecular warheads such as cytotoxic agents and radionuclides to tumors^1^. Meanwhile, engineered microbial and mammalian cells have provided therapeutic capabilities based on their ability to home to specific tissues^2,3^, express diagnostic^3–5^ or therapeutic^6–8^ proteins or carry out targeted cell killing^9,10^. However, both biomolecular and cell-based therapies are currently limited by a lack of mechanisms for external spatial and temporal control and a restricted repertoire of therapeutic actions^11,12^. Here we address these limitations by designing biomolecules and cells to act as seeds for acoustic inertial cavitation – the formation, growth and violent collapse of gas bubbles. This phenomenon is triggered remotely at specific locations using focused ultrasound, leading to strong local mechanical effects capable of killing target cells and actuating the release of genetically encoded therapeutic payloads.

Our approach takes advantage of gas vesicles (GVs), a unique class of genetically encoded gas-filled protein nanostructures that were recently introduced as reporter genes for high-frequency ultrasound and magnetic resonance imaging (MRI)^13–16^. GVs comprise amphiphilic protein shells with typical cylindrical widths of 45–250 nm and lengths of 100-600 nm^17^ that are permeable to gas but exclude water due to their hydrophobic interior surface (**Fig. 1a**). In nature, photosynthetic bacteria and archaea produce intracellular GVs as a means to achieve cellular buoyancy^18^. Recently it was discovered that GVs’ gas-filled interiors allow them to serve as contrast agents for high-frequency diagnostic ultrasound and MRI^13,14,16,19–22^. In addition, it was shown that engineered multi-gene clusters encoding GVs can be heterologously expressed in *Escherichia coli* and *Salmonella typhimurium*, two frequently used bacterial chasses for the development of cell-based diagnostic and therapeutic agents^15^. However, the use of GVs and GV-expressing cells as therapeutic agents has not been investigated.

**Figure 1:**
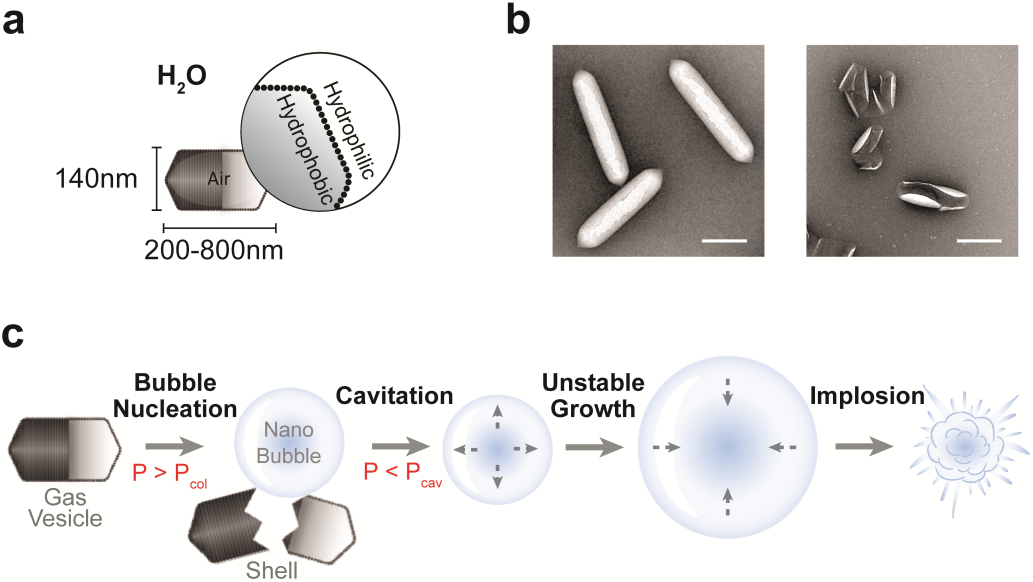
Gas vesicles as nuclei for inertial cavitation. **a**, Schematic drawing of a GV. The GV’s amphiphilic protein shell encloses a stable, gas-filled structure. **b**, Representative transmission electron microscopy (TEM) images of intact (left) and collapsed (right) *Anabaena flos-aquae* GVs. **c**, Proposed mechanism of GV-seeded cavitation. An ultrasound (US) pulse with a positive pressure higher than the critical collapse pressure, P_col_, collapses the GV, resulting in the release of a nanoscale air bubble. The released nanobubble undergoes cavitation if the peak negative pressure of the US pulse reaches below the critical cavitation pressure, P_cav_. Over several cycles, the nanobubble is converted into a micron-scale bubble, which can eventually undergo violent inertial cavitation. Scale bar represents 200 nm.

In this study, we hypothesized that at lower ultrasound frequencies GVs can serve as nuclei for the formation and cavitation of free bubbles, turning targeted and cell-expressed GVs into mechanical therapeutic warheads. We tested this fundamental hypothesis using acoustic spectroscopy and ultrafast optical microscopy, then asked whether GVs functionalized with targeting moieties can serve as ultrasound-triggered cellular disruptors. In addition, we assessed the ability of engineered therapeutic bacteria expressing GVs to be triggered with ultrasound to lyse and release molecular payloads. Finally, we tested the ability of GV-seeded cavitation to take place *in vivo* in a mouse tumor xenograft. These experiments provide a proof of principle for ultrasound-triggered inertial cavitation as a new capability in targeted molecular therapeutics and synthetic biology.

## RESULTS

### GVs act as seeds for bubble formation and cavitation

Our hypothesis that GVs can nucleate bubbles for inertial cavitation arises from the fact that GVs collapse under applied acoustic pressure (**Fig. 1b**), releasing the air contained inside them to the surrounding media. The ability of such collapse to take place at specific pressure thresholds, defined by GVs’ DNA sequence and protein composition, has been used for background-subtracted molecular imaging^13–15,19^. Under most conditions, gas molecules released from collapsed GVs are expected to form nanoscale bubbles, which should dissolve within milliseconds due to Laplace pressure^23^. However, we hypothesized that at ultrasound frequencies in the sub-MHz range, these free bubbles could also serve as seeds for cavitation, a process in which bubbles expand and contract during the negative and positive phases of sound waves, respectively, and can grow in size through rectified gas diffusion and coalescence^24^. Such processes are favored at lower ultrasound frequencies and higher peak negative pressures. In addition, bubbles can be stabilized by the presence of hydrophobic surfaces^25^, such as the exposed interior of collapsed GV shells (**Fig. 1b**). We envisioned that positive pressure above GVs’ critical collapse threshold would break open the GVs to release gas nanobubbles, and that negative pressure would then cause these bubbles to grow. At relatively low ultrasound amplitudes, the resulting bubbles would undergo stable cavitation – a sustained periodic oscillation of gas bubble size. At relatively high amplitudes, the bubbles would undergo rapid growth and violent collapse in a process known as inertial cavitation, unleashing powerful mechanical effects^26^ (**Fig. 1c**).

To test our hypothesis that GVs can seed bubble cavitation, we first performed acoustic measurements on GVs purified from *Anabaena flos-aquae*, looking for acoustic signatures of stable and inertial cavitation. While bubbles undergoing stable cavitation emit pressure waves at the transmitted frequency and its harmonics, those undergoing inertial cavitation emit pressure waves with broad spectral content. We performed acoustic spectroscopy on GV suspensions in a custom-built chamber with acoustically transparent polymer walls. Focused ultrasound (FUS) was applied with a focused single-element transducer, and emitted signals were recorded with an orthogonally positioned imaging transducer functioning as a passive cavitation detector (PCD) (**Fig. 2a**). Unless stated otherwise, GV were insonated at a frequency of 0.67 MHz throughout this study, and the PCD transducer was a 128-element linear array with a center frequency of 18 MHz. We chose this insonation frequency due to its common use in therapeutic ultrasound.

**Figure 2:**
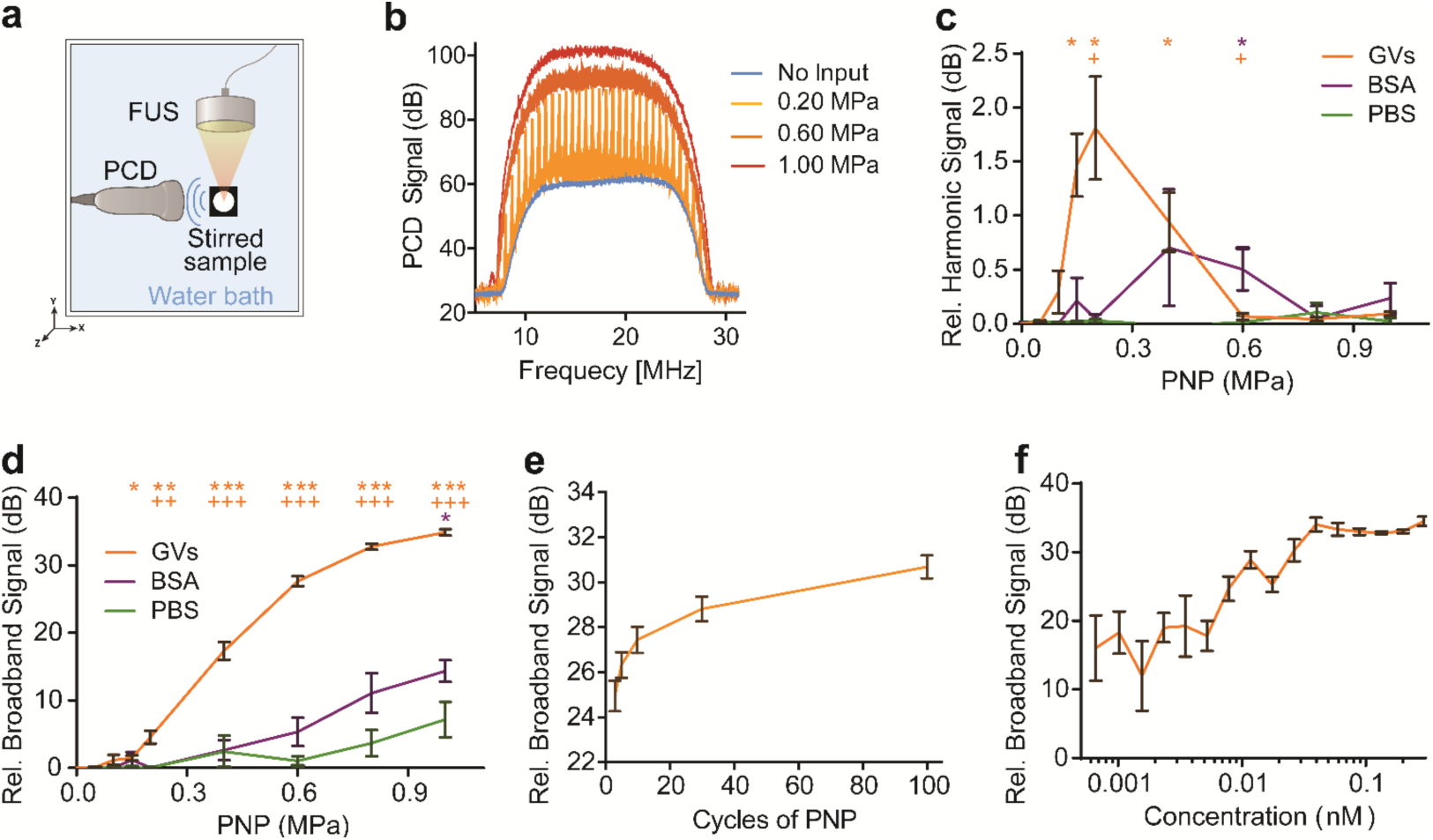
Purified GVs act as seeds for stable and inertial cavitation. **a**, Diagram of *in vitro* passive cavitation detection (PCD) setup used to measure the acoustic signatures of cavitation activity in response to focused ultrasound (FUS). **b**, Representative frequency spectra of backscattered signals from purified GVs (0.3 nM) insonated by a single US pulse at varying peak negative pressures (PNP), 30-cycles, 670 kHz. **C**, Mean harmonic signal from GVs (0.3 nM), bovine serum albumin (BSA, matched in mg/mL to GVs concentration), and PBS as a function of PNP (n = 16 for GVs and n = 8 for BSA and PBS). **d**, Mean broadband signal from GVs, BSA, and PBS as a function of PNP (n=16 for GVs, n = 8 for BSA, and PBS. statistical analysis: orange * - GVs vs. PBS, orange + - GVs vs. BSA, purple * - BSA vs. PBS). **e**, Average broadband measurements from GVs insonated with varying ultrasound pulse lengths (PNP = 0.6 MPa, n=12). **f**, Broadband signal from different concentrations of GVs insonated with a single 1.0 MPa pulse (n=5). Plots show mean ± SEM (c, d, e, f).

We observed both stable and inertial cavitation in GV suspensions exposed to FUS. Harmonic signals, indicative of stable cavitation, were clearly elicited when the GVs were insonated at 0.2 MPa peak negative pressure (PNP) (**Fig. 2b**, **c**), while broadband spectral emissions, characteristic of inertial cavitation, were observed at higher pressure levels (**Fig. 2b**, **d**). These values were significantly higher than those recorded from buffer or solutions of bovine serum albumin (BSA), which served as a protein control. Broadband emissions increased moderately with the pulse length (**Fig. 2e**), and pulses with only 3 peak negative cycles were sufficient to produce signals 25 ± 0.7 dB above the noise floor, consistent with cavitation studies on other samples^27,28^. As expected, the measured broadband signal increased with GV concentration until reaching a peak at 0.3 nM (**Fig. 1f**), above which acoustic shadowing interfered with measurement (**Supplementary Fig. 1**). These results suggest that GVs are able to serve as nuclei for inertial cavitation at 0.67 MHz.

Since GV are also used for ultrasound imaging, typically at frequencies of several MHz, we also measured cavitation responses at 3 MHz. At this frequency inertial cavitation required much higher pressures (**Supplementary Fig. 2a-b**), consistent with the lower efficiency of free bubble cavitation at higher frequencies, as well as the increased pressure required to collapse GVs at frequencies above the gas permeation rate of their protein shell^29^. This result affirms the ability of GVs to be imaged safely using typical diagnostic parameters^21,22^ while serving as seeds for inertial cavitation at lower frequencies.

To more directly visualize the process by which GVs nucleate the formation of cavitating bubbles, we imaged this process optically using an ultra-high frame rate camera, acquiring images at 5 million frames per second (**Fig. 3a**). The GVs were attached to acoustically transparent Mylar-bottomed dishes using biotin and streptavidin. Before insonation, we observed a dark pattern indicative of intact GVs (**Fig. 3b**), whose gas interiors scatter visible light^18,30^. After ultrasound was applied and reached sufficient amplitude, this dark pattern suddenly disappeared, revealing GV collapse (1.4 – 1.8 μs, **Fig. 3b** and **Supplementary Fig. 3**). 2.4 µs later, we observed dark bubbles forming and cavitating inside the field of view (**Fig. 3b**, **Supplementary Movie 1**), and continuing to grow in the following cycles (**Fig. 3d**). Meanwhile, control dishes with biotin coating alone failed to show significant cavitation (**Fig. 3c**). We further analyzed the videos to track the temporal relationship between GV collapse and bubble cavitation. After forming, bubbles grew and shrank at the frequency of the ultrasound waves (**Fig. 3d**). By comparing the phase of the wave at which GVs disappear with the phase of maximal bubble growth rate, we could confirm that GVs collapse at the positive pressure peak, while maximal growth of the resulting bubbles occurs at the negative peak of the ultrasound cycle, π apart in phase (**Fig. 3e**), and bubble size peaks at 3/2 π, at the conclusion of rarefaction (**Fig. 3f**). Similar results were seen across bubbles (**Fig. 3g**). This data confirms the ability of GVs to nucleate bubble cavitation and supports the mechanistic model depicted in in **Fig. 1c**.

**Figure 3:**
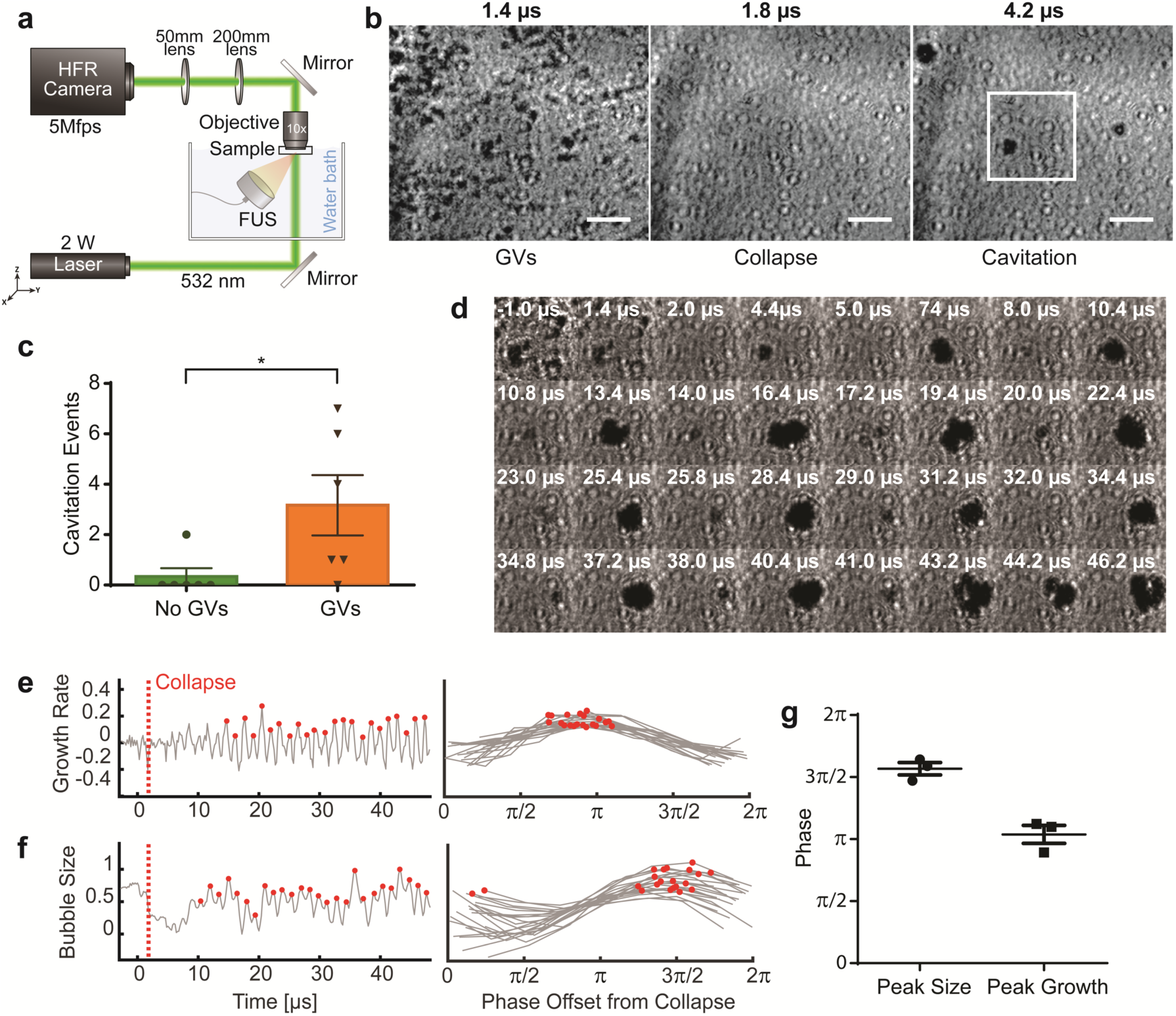
Ultrafast optical imaging of GV-seeded bubble formation and cavitation. **a**, Schematic drawing of the high frame rate (HFR) camera setup enabling GV cavitation imaging at a frame rate of 5 MHz. **b**, HFR camera images immediately before GV collapse (left), immediately after GV collapse (middle), and after the formation of bubbles (right). **c**, Number of unique cavitation loci in biotinylated dishes with and without GVs, upon insonation with a single 1.4 MPa burst (p= 0.0411, n=6). FUS pulses with 30 cycles at 670 kHz were used unless otherwise stated. **d**, Representative high-speed camera frames showing every other maximum and minimum of bubble cavitation, preceded by GV collapse. **e**, Bubble growth rate, quantified as the temporal derivative of the normalized average inverted pixel intensity in (d) (left). The plot on the right shows each maximum in the growth rate aligned to the phase offset from the time of GV collapse. **f**, Bubble size and phase offset from GV collapse analyzed from HFR images as in (e). **g**, Average phase offset for peak size and peak growth rate for three different regions of interest. Plots show mean ± SEM (c, g). Scale bar represents 20 µm (b).

### Receptor-targeted GVs serve as acoustically detonated cellular disruptors

After establishing that GVs can nucleate microscale bubbles, we investigated two applications of biomolecular cavitation: acoustically detonated killing of tumor cells with receptor-targeted GVs and triggered cavitation of intracellular GVs in engineered bacteria, leading to cell lysis and release of a protein payload. To test the first application, we prepared GVs genetically engineered to display an RGD peptide on the C-terminus of their outer shell protein, GvpC, thereby targeting them to αVβ_3_ integrin receptors commonly overexpressed in tumors^31^. We incubated these nanostructures with U87 glioblastoma cells cultured on Mylar film (**Fig. 4**, **a**-**b**). For visualization, the GVs were also chemically labeled with Alexa Fluor 488. To monitor cellular disruption, the media was supplemented with propidium iodide (PI), a membrane-impermeable fluorophore that becomes fluorescent upon entering disrupted cells and intercalating with nucleic acids. Prior to ultrasound exposure, there was negligible PI signal. However, after insonation for 10 seconds, we observed gradual PI uptake in many cells throughout the field of view (**Fig. 4**, **c-d**), as quantified relative to PI uptake after saponin treatment (**Fig. 4e**). Ultrasound alone in the absence of GVs did not result in significant PI uptake.

**Figure 4:**
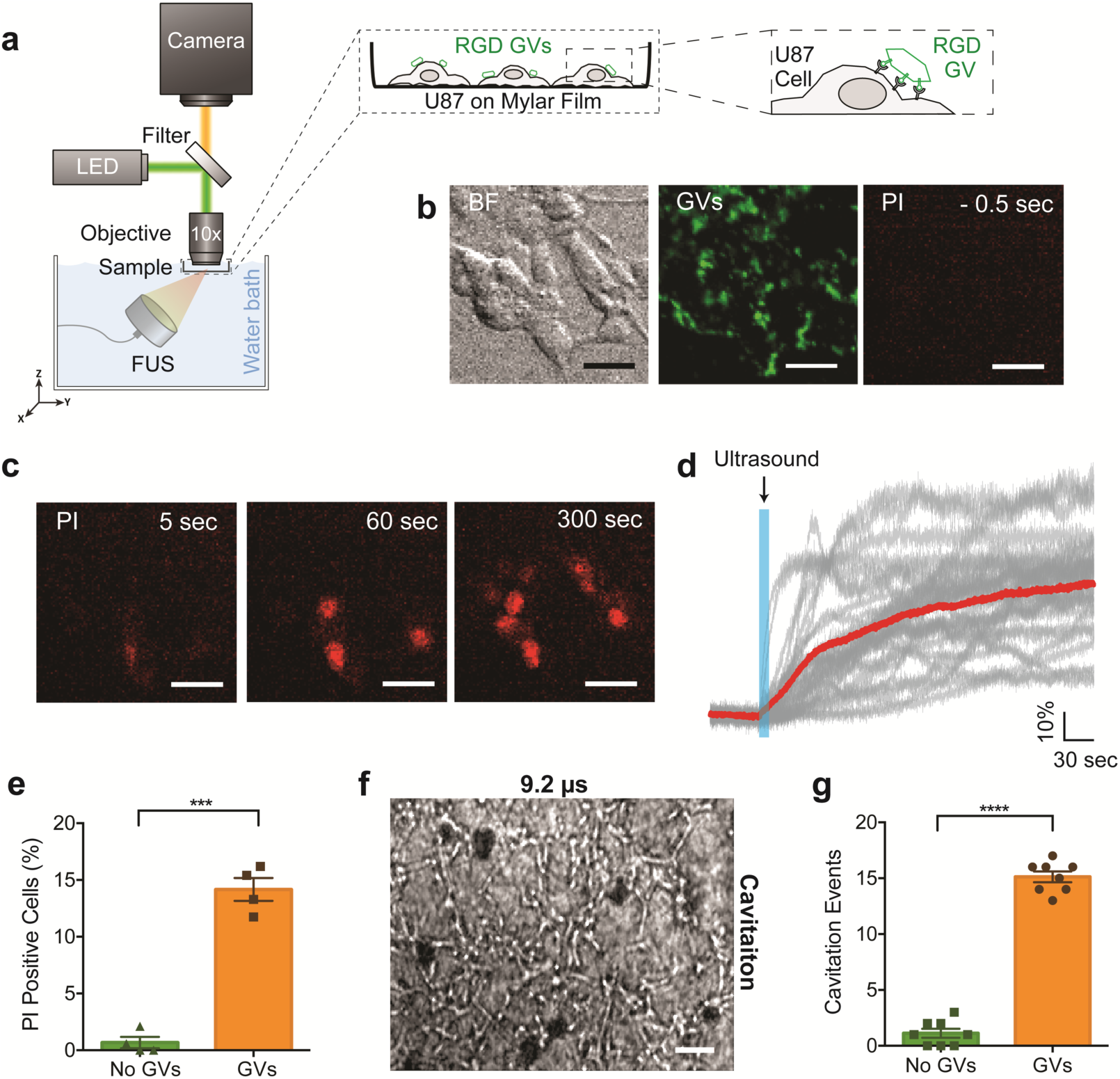
Molecularly-targeted GVs serve as ultrasound-triggered disruptors of mammalian cells. **a**, Schematic drawing of the fluorescent microscopy setup used for imaging of GV-mediated cell disruption. RGD-functionalized GVs were attached to U87 cells grown on Mylar-bottomed dishes. **b**, Bright field (BF) image of U87 cells, fluorescence images of GVs (green) and propidium iodide (PI, red), before the application of ultrasound. **c**, PI fluorescence 5, 60, or 300 sec after ultrasound exposure. **d**, Change in PI signal measured from individual cells (gray) and the average (red) before, during and after FUS application (represented by blue shading). **e**, Percentage of PI-positive cells following FUS exposure with and without GV attachment. (P = 0.0002 using a two-sided heteroscedastic t-test, n=4) **f**, HFR camera image showing the formation of bubbles during ultrasound application to cells treated with GVs. **g**, Number of unique cavitation loci observed in dishes containing U87 cells with and without GVs. (P < 0.0001, n=8). Bar plots show mean ± SEM (e, g). Scale bars represent 20 µm (b, c, f).

To directly visualize the effect of ultrasound on GVs attached to U87 cells, we imaged the cells at 5 million frames per second. After the collapse of cell-attached GVs (**Supplementary Fig. 4**, **a-b**) we observed bubble formation and cavitation (**Fig. 4f**, **Supplementary Fig. 4c**, **Supplementary Movie 2**). The number of cavitation events in cells targeted with GVs was significantly greater than the number of random cavitation events in untreated cells (**Fig. 4g**). The spatial heterogeneity of cavitation in the high-speed camera experiment (**Fig. 4f**, **Supplementary Fig. 4c**) was consistent with the clustering of PI-positive cells next to each other (**Fig. 4c**). Taken together, these results suggest that GVs can be used as targeted, acoustically triggered mechanical warheads for cellular disruption.

### Genetically expressed GVs enable cells to undergo inertial cavitation

In addition to their use as purified agents, GVs can be expressed inside engineered cells, as shown recently in *E. coli Nissle 1917* and *S. typhimurium* using a hybrid GV-encoding gene cluster^15^. We hypothesized that cells expressing GVs could be triggered to undergo intracellular bubble formation and cavitation under low frequency ultrasound, resulting in cellular lysis and the release of a co-expressed protein payload (**Fig. 5a**). To test this concept, we engineered a 14-gene operon combining GV-encoding genes GvpA–GvpU from the ARG1 gene cluster^15^ with a gene encoding the luminescent NanoLuc protein as a model releasable payload (**Fig. 5b**). This construct was transformed into *S. typhimurium* SL1344, a strain used in several experimental bacterial therapies^2,3^. Since it was previously shown that ARG1 GVs have a relatively high collapse pressure^15^ we used a lower ultrasound frequency of 300 kHz in these experiments to ensure efficient collapse of the heterologously expressed GVs at achievable pressure levels.

**Figure 5:**
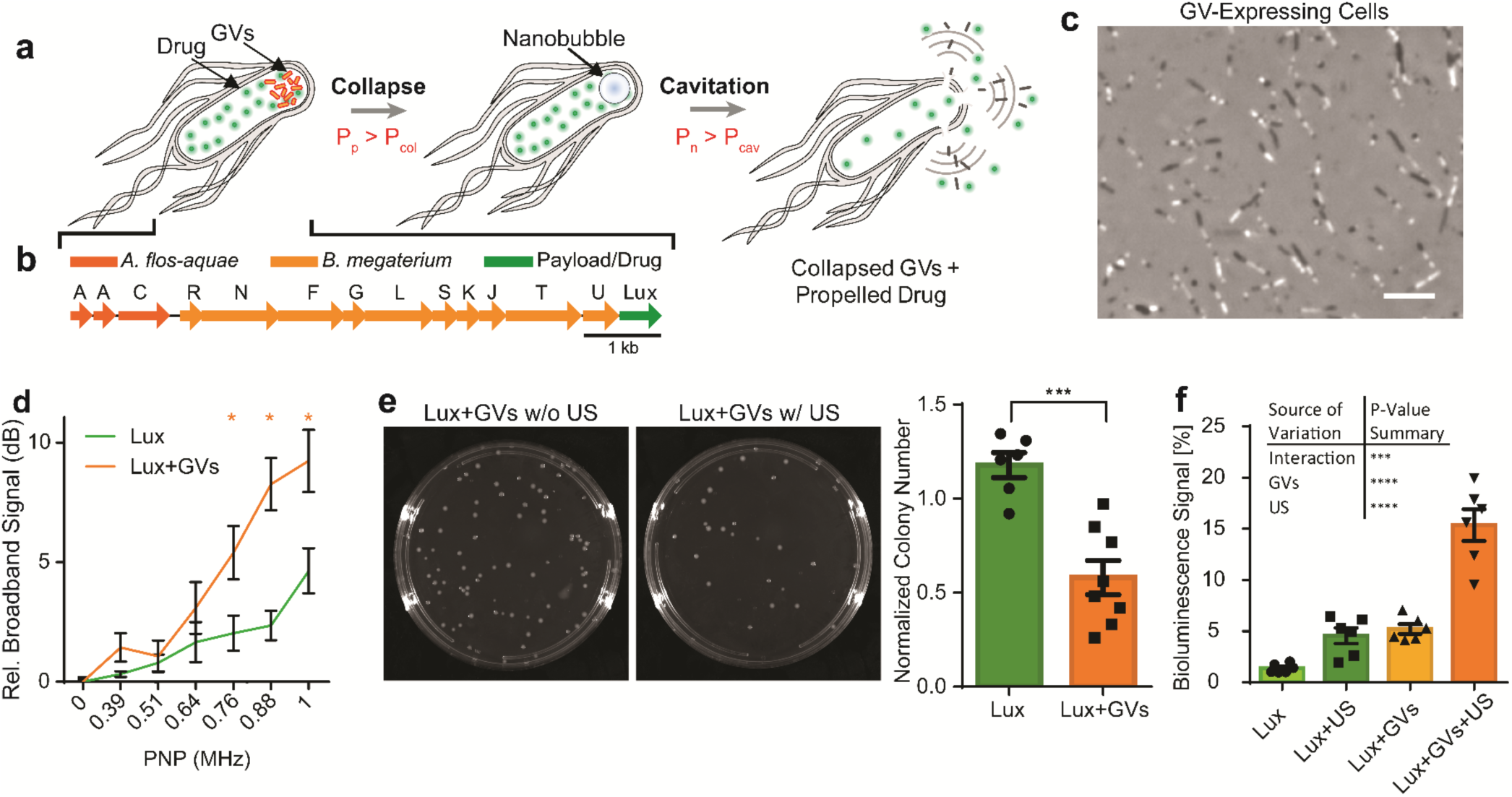
GVs as genetically encoded seeds for cellular inertial cavitation and payload release. **a**, Proposed mechanism of intra-cellular GV-seeded cavitation and cell disruption. The collapse of GVs inside the cell creates a nanobubble, which cavitates, disrupts the cell membrane and releases cell contents, including an engineered payload. **b**, Genetic construct combining a hybrid gas vesicle gene cluster from *A. flos-aquae* and *B. megaterium* with the a NanoLuc luciferase (Lux) payload. **c**, Phase contrast microscopy image of *S. typhimurium* cells expressing the construct in (b). GVs appear inside the cells as white inclusions, while the rest of the cells appear black. **d**, Mean broadband emissions from NanoLuc-expressing cells (Lux, negative control) and cells co-expressing GVs and NanoLuc, at various pressure levels (p < 0.05, n = 8) **e**, Representative agar plates and average colony counts for Lux and Lux+GV cells exposed to ultrasound, normalized by the number of colonies from non-exposed control samples (p = 0.0002, n = 8 for Lux+GV and n = 6 for Lux). **f**, Bioluminescent signal in the media surrounding Lux or Lux+GV cells with and without exposure to ultrasound. The signal from each sample is normalized by the signal measured from the same cells after chemical lysis (two-way ANOVA analysis: percentage of variation attributed to the interaction is 9.6% with p=0.001, n = 6). Scale bar represents 15 μm (c).

To test whether the GV-expressing bacteria (**Fig. 5c**) could serve as sources of inertial cavitation, we measured broadband acoustic emissions from cell suspensions exposed to FUS. In response to ultrasound pulses, the GV-expressing *S. typhimurium* emitted a high level of broadband signals, increasing with the peak negative pressure (**Fig. 5d**). Similar activity was not observed in control cells expressing just NanoLuc.

Next, we examined the lysis of GV-expressing cells in response to focused ultrasound and measured the release of their co-expressed protein payload. In an assay comparing the number of colonies formed on agar plates by cells that were exposed to ultrasound to cells that were not, we found that significantly fewer colonies were produced by GV-expressing *S. typhimurium* cells following ultrasound exposure, compared to the equivalent experiment with NanoLuc controls (**Fig. 5e**). The bioluminescence of the media, corresponding to payload release, was also significantly elevated in GV-expressing cells exposed to ultrasound compared to controls (**Fig. 5f**). Taken together, these results suggest that GV-expressing engineered cells can serve as ultrasound-triggered cellular “explosives”, releasing proteins into their surroundings in response to a remote trigger.

### GVs seed inertial cavitation in vivo

After establishing GVs as biomolecular and cellular cavitation nuclei *in vitro* and *in cellulo*, we tested the ability of GVs to nucleate cavitation *in vivo* in a mouse tumor xenograft. We developed a 3D-printed holder to co-align the foci of both the focused transducer and the imaging transducer at the center of the tumor (**Fig. 6a**), with a needle guide incorporated to facilitate precise injections into the tumor core. This setup enabled us to perform ultrasound-guided ultrasound therapy experiments, in which we acquired images of the GVs inside the tumor and performed FUS treatment and PCD measurements at the injection site. Adult BALB/c mice were injected in the flanks with two MC26 tumors, one in each side. After the tumors reached diameters of 6-10 mm (**Fig. 6b**, left), we performed ultrasound-guided GV injections and FUS treatment. We injected one tumor in each mouse with purified GVs and the contralateral tumor with saline as a vehicle control. To facilitate GV imaging, we used GVs in which GvpC has been removed^19^, allowing them to produce non-linear ultrasound contrast easily distinguishable from background using an amplitude modulation pulse sequence^22^ (**Fig. 6b**, middle). PCD measurements acquired during FUS application, performed before and after the injection of saline or GVs, revealed strong broadband signals representing inertial cavitation only from GV-injected tumors (**Fig. 6**, **c-d**). After FUS exposure, the image contrast from injected GVs disappeared, providing confirmation of their collapse (**Fig. 6b**, right). These results demonstrate that gas vesicles can serve as biomolecular cavitation nuclei in a disease-relevant tissue *in vivo*.

**Figure 6:**
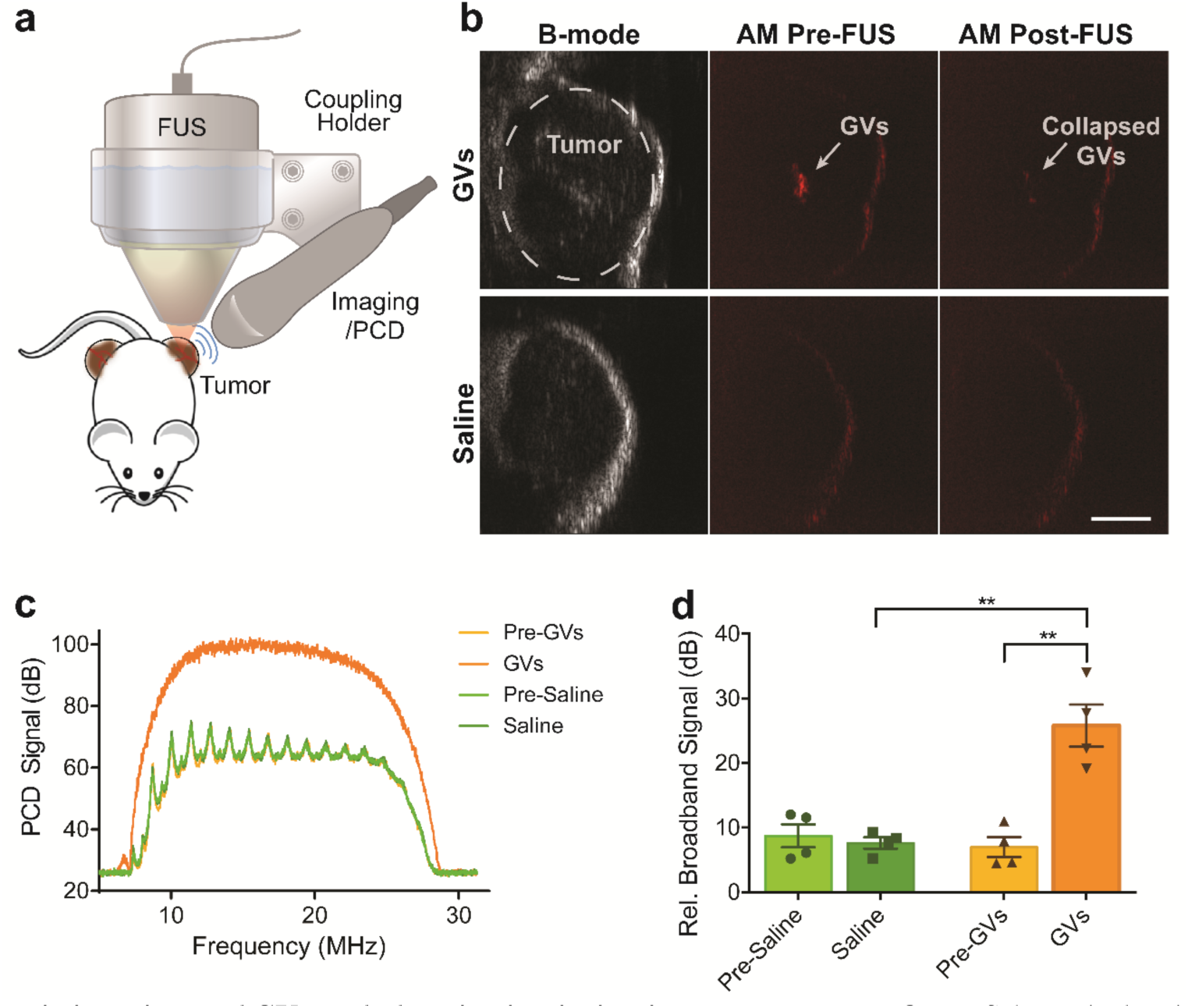
Theranosic imaging and GV-seeded cavitation in *in vivo* tumor xenografts. **a**, Schematic drawing of *in vivo* FUS and imaging/PCD setup. A 670 kHz FUS transducer and a 18 MHz imaging linear array transducer are aligned using a 3D-printed cone holder. Balb/c mice harbor bilateral MC26 subcutaneous hind-limb tumor xenografts. **b**, Ultrasound images of tumors injected with engineered nonlinear GVs or saline (sham control). The grayscale B-mode images present the anatomy of the tumor, while amplitude modulation (AM) imaging shows GV-selective nonlinear contrast, which disappears after FUS application. **c**, Representative PCD spectra measured before and after injection of purified GVs (2.7 nM) or saline into tumors. Single 1.0 MPa, 30-cycle long pulses were transmitted from a 670 kHz FUS transducer. **d**, Relative broadband signal measured from pre-and post-GV and saline injections into tumors (p=0.002, n=4). Bar plots show mean ± SEM (d). Scale bar is 3 mm (b).

## DISCUSSION

This work establishes GVs as the first genetically encodable nuclei for inertial cavitation, bringing together the unique capabilities of targeted therapeutics, synthetic biology, focused ultrasound and bubble mechanics. Our fundamental finding that GVs are capable of nucleating cavitation both in their purified form and inside genetically engineered cells will enable two distinct types of future applications. First, purified GVs targeted to cells such as tumors via specific surface markers can serve as ultrasound-triggered disruptors of the plasma membrane, causing cell death and making the interior of the target cells accessible to synergistic drug molecules. The development of this application will benefit from GVs’ fundamental physical stablitiy^13^, the tunability of their size and shape, and the ability of their surfaces to be functionalized via genetic fusion or chemical methods.

Second, GVs expressed by engineered cells will provide a new capability for therapeutic synthetic biology. Compared to conventional drugs, cell-based therapeutic agents such as engineered bacteria have a broader range of functionality based on their ability to migrate to diseased tissues such as tumors^2,3^ or the GI tract^7^, recognize specific molecular signals, proliferate, and produce therapeutic biomolecules. Giving such agents the ability to express their own GVs as intracellular cavitation nuclei turns them into ultrasound-triggered cellular explosives, whose remote actuation results in the release of intracellular therapeutic payloads. In addition, the accompanying bacterial lysis may help activate the immune system^32^, thereby enhancing the efficacy of co-administered immunotherapy^2,33^. Furthermore, the ability of focused ultrasound to specify where and when in the body these effects take place could help overcome off-target toxicity arising from cell migration to organs such as the spleen, liver, and kidney^34,35^. Additionally, the conditional production of GVs in response to specific biochemical signals could add another layer of control.

The development of GVs as therapeutic agents will further benefit from their ability to be visualized with non-invasive imaging modalities such as ultrasound and magnetic resonance imaging, allowing theranostic treatment planning and verification. The unique contrast produced by GVs can be combined with other imaging modes providing information about tissue features such as vascularization, oxygenation, and elasticity^36–38^. In this context, one of the important findings of this work is the clear distinction between the ultrasound frequencies and pressure levels needed for GV cavitation compared to those used for imaging. In addition, the disappearance of GV image contrast after cavitation provides a means to confirm the spatial targeting of therapeutic activation.

The proof-of-concept experiments presented in this work establish the need for essential follow-up studies to further understand and apply the biomolecular acoustic cavitation paradigm. Higher-resolution optical imaging is needed to more fully visualize GV collapse, nanoscale bubble formation, growth and cavitation. Currently only GV collapse and micron-size bubble formation could be seen, and it was not possible to quantify the efficiency of each step in this overall process. Additional fundamental studies are also needed to characterize the impact of GV clustering at the cellular surface or interior, and of the surrounding biochemical composition, on bubble nucleation. Based on the insights from such studies, GVs and GV-expressing cells can be further engineered to increase their capabilities as cavitation nuclei. For example, genetic engineering allows the tuning of GV particle properties such as critical collapse pressures, dimensions, and surface chemistry^19,31^. GV-expressing cells can also be tuned to adjust expression levels for GVs and therapeutic payloads, the timing or conditional triggering of expression, and other cellular properties such as wall and membrane composition, which may influence cavitation and lysis. In addition, ongoing efforts to express GVs in mammalian cells will greatly expand the range of cellular sonotherapeutics. Finally, building on the fundamental findings and basic proof-of-concept demonstrations performed in this study, future work must test specific therapeutic applications in more sophisticated disease models. We expect the discovery of genetically encoded nuclei for inertial cavitation to produce an explosion of future studies and applications.

## MATERIALS AND METHODS

### Chemicals

All chemicals were purchased from Sigma Aldrich unless otherwise noted.

### GV purification, modification, and quantification

*Anabaena flos-aquae* GVs were produced and purified as described previously^31^. As a quality assurance step, OD measurements were performed for each sample at 0-12 bar using an echoVis Vis-NIR light source coupled with an STS-VIS spectrometer (Ocean Optics), and a 176.700-QS sample chamber (Hellma Analytics). GvpC was removed from GVs used in the *in vivo* experiment to facilitate their ultrasound detection. GvpC was removed and replaced by an engineered, recombinantly expressed, protein, GvpC-RGD, for experiments in which GVs were attached to U87 cells^19^. This GvpC variant (GvpC-R3) was also shorter than the wild type protein, resulting in GV collapse at lower pressure levels^19^. Prior to experiments, GV concentrations were measured using a spectrophotometer (Thermo Scientific NanoDrop^TM^ ND-1000) at 500nm.

### *In vitro* passive cavitation detection

The *in vitro* setup (**Fig. 2a**) was aligned in a three-step process. First, an L22-14v 128-element Verasonics imaging probe was positioned such that its focus, set to a depth of 8 mm, was at the center of a 1 cm × 1 cm 3D-printed chamber. Then, an optic fiber hydrophone (Precision Acoustics) was positioned at the center of the chamber using B-mode imaging. Finally, a 670 kHz (Precision Acoustics) or 3 MHz FUS transducer (Ultrasonic S-Lab) was mounted on a computer-controlled 3D translatable stage (Velmex) orthogonally to the Verasonics probe. A MATLAB program automatically scanned and aligned the transducer’s focus at the center of the chamber, according to the feedback from the hydrophone.

A solution of purified GVs (OD_500_ = 0.5, or 0.3 nM, unless stated differently) or GV-expressing bacteria (OD_600_ = 1) was gently pipetted into the 3D printed chamber to minimize the introduction of bubbles. Purified GVs solutions had been prepared several days in advance to allow natural degassing to atmospheric conditions. The Verasonics scanner was programmed to function in zero-amplitude transmit so it could be used as a passive cavitation detector. A MATLAB script was written to synchronize the acquisition of GV signals by triggering the FUS burst and accounting for the propagation time of the insonating wave to the focus. The GV solution was stirred gently during acquisition using a micro stir bar and magnetic stirrer (Thermo Scientific).

To compensate for the slow rise time of the Precision Acoustics transducer, a custom waveform with 33 cycles was created such that the first 4 cycles had 1.5x the amplitude as the last 29 cycles; this waveform induced our FUS to reach the desired peak negative pressure (PNP) output at the 4^th^ cycle. In experiments where the effect of the number of cycles on cavitation was investigated, a waveform with 3 cycles at 2x the amplitude of the remaining cycles was used so that the desired PNP was reached at the 3^rd^ negative peak.

### Mammalian cells experiments

Glass bottom 35-mm petri dishes (Matsunami) were modified to enable U87 cell culture and subsequent ultrasound application. The glass was removed using a glass cutter and Mylar film (Chemplex, 2.5 μM thickness) was fixed over the hole via a polydimethylsiloxane elastomer (Sylgard 184 silicone, Dow Corning). After curing for 24 hours at 40°C, these dishes were sterilized using UV light. U87 cells were plated on the Mylar film dish and incubated for 2 days at 37°C in 2mL of DMEM media. In order to facilitate the attachment of RGD-GVs to the cells membrane, the GVs were re-suspended in fresh DMEM (final concentration of 0.5 OD_500_) and added to the center of the dishes. The center of each dish was sealed using an 18-mm round cover glass and kept inverted for 2 hours at room temperature. Then, cells were washed and recovered with fresh DMEM. Finally, 10 μg/ml propidium iodide (PI, Invitrogen) was added to the medium just before the ultrasound experiment.

In this experiment, the cells were insonated using a 670 kHz transducer (Precision Acoustics) that was positioned in the water tank at an angle of 20° relative to the water surface, to minimize standing waves (**Fig. 4a**). The ultrasound transducer was aligned to the microscope using a hydrophone. The cells were insonated for 10 seconds with PNP = 1.5 MPa ultrasound pulses at a 2 ms pulse repetition rate. Fluorescence recording began 1 minute before insonation, continued throughout ultrasound exposure, and ended 10 minutes after insonation. Fluorescence signals were collected using a 10x immersion objective (Olympus, NA 0.3), and a sCMOS camera (Zyla 5.5, Andor) at a 10 Hz frame rate. After this acquisition, we used saponin (Sigma, 100 μg/ml) to perforate all cell membranes, and the resulting image was used as a mask for cell body detection. The fluorescence images were processed using NeuroCa^39^ to extract fluorescent signals from individual cells. Cells were defined as PI-positive if signal intensity increased by more than 2% following ultrasound application.

### High frame rate camera imaging experiments

We assembled a high-speed microscopy setup capable of directly capturing GV collapse and bubble cavitation events (**Fig. 3a**). Our setup used a 2-W 532-nm laser (CNI, MLL-F- 532-2W) controlled by an optical beam shutter (Thorlabs SH05, KSC101). Right angle prism mirrors directed the laser light through a water bath and into a sample dish containing the imaged samples. For imaging purified GVs, the GVs were biotinylated by incubating them for 1 hour with a 10,000-fold molar excess of sulfo-NHS-biotin (Thermo Fisher Scientific). Then, the non-attached biotin was removed by dialysis in PBS. The Mylar dishes were prepared by first treating them with UV light, then incubating them with 180 µL of 0.1 mg/mL poly-D-lysine hydrobromide. The dishes were then incubated with sulfo-NHS-biotin (Thermo Fisher Scientific, 180 µL, 2mM) for 1 hour. After washing away free biotin with PBS, dishes were incubated with streptavidin (G-Biosciences, 180 µL, 7.35µM) for 1 hour and washed again to remove unbound streptavidin. Finally, the biotinylated GVs were attached to the dishes. After adding 180 µL of GVs at OD_500_ = 2 to each dish, the center of each dish was sealed using an 18 mm round cover glass and kept inverted for 2 hours at room temperature. Then, excess GVs were washed away with PBS. All wash steps were repeated at least 3 times. For imaging cells, the same dishes and cell culture processes described in the previous section were used, except with a final GV concentration of OD_500_ = 2. Dishes containing GVs or cells were positioned above a water tank and aligned with the transducer focus as described above. A 10x water immersion Plan Fluor objective (Olympus, NA 0.3) was lowered into the solution in the dish. A series of prism mirrors and converging lenses with focal lengths of 200 mm and 50 mm delivered the image into a Shimadzu HPV-X2 camera, which acquired 256 images (**Supplementary Fig. 3**) over 51.2 µs, at a sampling rate of 5 million frames per second. To account for acoustic propagation through water, the camera was externally triggered to begin acquisition 40 µs after the start of the ultrasound pulse. A single pulse with 30 cycles and PNP = 1.4 MPa was used to insonate the sample in these experiments.

### Bacterial expression and experiments

GV-expressing *Salmonella typhimurium* were produced by transforming cells with a plasmid encoding an engineered genetic construct comprising a GV operon ^15,31^ and a NanoLuc luciferase. Cells transformed with a NanoLuc-only plasmid were used as controls. Constructs were assembled using Gibson cloning. The genetic constructs were cloned into the pTD103 plasmid (gift from J. Hasty), with expression driven by a luxI promoter upon induction with 3nM N-(β-ketocaproyl)-l-homoserine lactone (AHL). The cells were cultured for 24 hours at 30 °C after induction, then centrifugated for 4 hours at 150 × g and 4 °C to enrich for buoyant cells. Samples of cells used in PCD experiments were stored for two days at 4 °C before these experiments and were always gently pipetted so as to minimize media gassing and bubble formation. Cells used for the NanoLuc release experiment were washed four times by 2 hours of centrifugation at 150 × g and 4 °C to remove any NanoLuc molecules that may be present in the media prior to the experiment.

PCD recordings from GV-expressing cells were performed using the same setup and protocol used in PCD recordings from purified GVs, with cells at a concentration of OD_600_ = 1. The same experimental setup was also used in the bacteria lysis and the payload release experiments; however, in these experiments, samples were insonated for 30 seconds at 300 kHz, PNP = 1 MPa, and a pulse repetition interval of 2 ms. In addition, to place the cell samples at the main lobe of the ultrasound beam, the sample chamber was filled with 1% agar, leaving a 4 mm diameter empty well, located in its center. Cell samples were loaded into this well at a concentration of OD_600_ = 2 and a volume of 50 mL, and covered with paraffin oil. For colony counts, the cells were plated on agar plates with kanamycin. Plates were imaged using a ChemiDoc Gel Imaging System (Bio-Rad) using a white epi-illumination protocol. Then, the colonies were counted to determine total colony forming units. In the payload release experiments, the solution was aspirated from the chamber after exposure to ultrasound, pipetted into 100 kDa Spin-X(R) UF concentrators (Corning) and centrifuged at 300 × g for 30 minutes to separate the supernatant fluid from the pellet and the buoyant cells. Then, the NanoLuc signal was measured using a Nano-Glo Luciferase assay kit (Promega) and a plate reader system (molecular devices). Full chemical lysis of cells using SoluLyse^TM^ Protein Extraction Reagent (Genlantis) was used as positive control.

### *In vivo* passive cavitation detection

All *in vivo* experiments were performed on BALB/c female mice, under a protocol approved by the Institutional Animal Care and Use Committee of the California Institute of Technology. No randomization or blinding were necessary in this study. Mice were anaesthetized using 1–2.5% isoflurane during all the injection and imaging procedures. The MC26 colorectal cancer cell line were maintained per standard cell culture techniques. Four female BALB/c mice, aged 8 weeks, were given subcutaneous inoculations of 5 × 10^6^ MC26 cells into both right and left hind flanks. Tumors were monitored and permitted to grow to a diameter of 6-10 mm over 10-20 days.

A 3D-printed theranostic holder co-aligned a 0.67 MHz FUS transducer (Precision Acoustics) and an L22-14v 128-element imaging probe (Verasonics) by fixing the FUS focus at 12 mm along the imaging plane (**Fig. 6a**). The holder places the FUS cone facing downwards and the imaging probe at approximately 30° from the vertical. A 3D-printed needle guide was mounted to the side of the cone such that the tip of a 1-inch 30-gauge injection needle also intersected the focus. The holder was mounted on a manually-controlled 3D positioner.

Mice were anaesthetized, maintained at 37 °C on a heating pad, depilated over the tumor region, and positioned with tumor facing directly upwards. Prior to the experiment, the ultrasound gel was centrifugated at 2000 × g for 10 minutes to remove bubbles, heated to 37 °C, and then carefully applied to couple the cone and probe to the tumor. B-mode anatomical imaging was used to confirm the absence of bubbles in gel application and to position the center of the tumor at an axial depth of 12 mm. B-mode and amplitude modulation (AM)^22^ images of the tumor were saved pre- and post-insonation. Insonation comprised a single 30-cycle, PNP = 1 MPa burst. The same PCD script was used as for *in vitro* PCD acquisitions.

As part of the experiment, 20 μl of OD_500_ = 4.5 GVs (with GvpC removed) or saline were infused directly into the tumor at a flow rate of 10?μl min^-1^ via a Genie Touch syringe pump through PE10 catheter tubing and a 30-gauge needle (BD). Injection at the focal zone was confirmed via B-mode imaging by locating the needle tip, and AM and B-mode images were recorded pre-and post-insonation. PCD measurements were performed during each insonation.

### Passive cavitation detection data processing

The acoustic emissions acquired by the PCD were sampled by the Verasonics scanner at 62.5 MHz and processed using a MATLAB (2017b, Mathworks) script. Single channel, 8192 - point FFT frequency spectrum estimations of the RF recordings from the 128 transducer elements were averaged to produce each PCD frequency spectrum estimation. To calculate the average amount of stable cavitation, an acquisition with clear harmonic response was used to manually select and save the peak harmonic frequencies (see **Supplementary Table 1**). Then, a trend curve was fitted to each spectral estimation using the MATLAB Curve Fitting tool. The smoothing parameter was chosen such that the resulting curves included only the broadband signal and not any harmonic peaks. This smoothened curve was subtracted from the original frequency spectrum, and the resulting flattened spectrum with harmonic peaks was integrated over the peaks, each with a bandwidth of 191 kHz for the 670 kHz measurements, or 610 kHz for 3 MHz measurements (so as to include the full harmonic peak). Integrals were performed using trapezoidal sums, then the integral was divided by the product of the number of peaks and the peak bandwidth.

To calculate the average amount of broadband emission, baseline noise at PNP = 0 MPa was subtracted from each reading. Then, the flattened spectrum with harmonic peaks was subtracted. The remaining spectrum was integrated between 7.4 MHz and 28.6 MHz, which marked the beginning and end of baseline noise, then divided by 21.2 MHz.

### Statistical analysis

For statistical significance testing, we used two-sided heteroscedastic t-tests with a significance level of type I error set at 0.05 for rejecting the null hypothesis, unless mentioned otherwise. A Wilcoxon rank sum test was used in high-speed camera experiments that included a control group that was found to be non-Gaussian using a Lilliefors test (P < 0.001). A two-way ANOVA test was used in the payload release experiment. Sample sizes for all experiments, including animal experiments, were chosen on the basis of preliminary experiments to be adequate for statistical analysis.

### Data and code availability

MATLAB codes are available from the corresponding author upon reasonable request. Plasmids sequences are included in Supplementary Information (**Supplementary Table 2**), and plasmids will be made available through Addgene upon publication. All other materials and data are available from the corresponding author upon reasonable request.

## ACKNOWLEDGEMENTS

The authors thank Dan Piraner, Anupama Lakshmanan, Arash Farhadi, and Pradeep Ramesh for helpful discussions. We thank Arash Farhadi for help with the GvpC-RGD variant and Hunter Davis for input on HFR imaging optics. We thank Maayan Harel (www.maayanillustration.com) for the illustrations in this paper. We thank Alasdair McDowall for help with electron microscopy. This project was supported by the David and Lucile Packard Fellowship for Science and Engineering (MGS) and the Heritage Medical Research Institute (MGS). AB-Z is supported by the Marie Skłodowska-Curie Fellowship and the Lester Deutsch Fellowship. AN was supported by the Amgen Scholars program. MA is supported by the NSF Graduate Research Fellowship and the P.D. Soros Fellowship. D Maresca is supported by the Human Frontiers Science Program Cross-Disciplinary Fellowship.

## AUTHOR CONTRIBUTIONS

AB-Z and MGS conceived the study. AB-Z, AN, D Maresca, DRM, and SY designed, planned and conducted *in vitro* experiments. AB-Z, AN, and AL-G designed, planned and conducted *in vivo* experiments. AB-Z edited the gene circuits with MA guidance. AB-Z, AN, DRM, SY, and D Maresca analyzed the data. D Malounda prepared the purified GVs. All authors discussed the results. AB-Z, AN, and MGS wrote the manuscript with input from all the authors. All the authors have given their approval for the final version of the manuscript. MGS supervised the research.

## References

1. Milenic, D. E., Brady, E. D. & Brechbiel, M. W. Antibody-Targeted Radiation Cancer Therapy. 3, (2004).

2. Din, M. O. et al. Synchronized cycles of bacterial lysis for in vivo delivery. Nature 536, 81–85 (2016).

3. Danino, T. et al. Programmable probiotics for non-invasive urinary detection of cancer. Sci. Transl. Med. 7, 1–36 (2015).

4. Kotula, J. W. et al. Programmable bacteria detect and record an environmental signal in the mammalian gut. Proc. Natl. Acad. Sci. 111, 4838–4843 (2014).

5. Archer, E. J., Robinson, A. B. & Süel, G. M. Engineered E. coli that detect and respond to gut inflammation through nitric oxide sensing. ACS Synth. Biol. 1, 451–457 (2012).

6. Claesen, J. & Fischbach, M. A. Synthetic microbes as drug delivery systems. ACS Synth. Biol. 4, 358–364 (2015).

7. Daniel, C., Roussel, Y., Kleerebezem, M. & Pot, B. Recombinant lactic acid bacteria as mucosal biotherapeutic agents. Trends Biotechnol. 29, 499–508 (2011).

8. Steidler, L. et al. Treatment of Murine Colitis by Lactococcus Secreting Interleukin-10. Adv. Sci. 289, 1352–1355 (2011).

9. Davila, M. L. et al. Efficacy and Toxicity Management of 19-28z CAR T Cell Therapy in B Cell Acute Lymphoblastic Leukemia. 6, (2014).

10. Jackson, H. J., Rafiq, S. & Brentjens, R. J. Driving CAR T - cells forward. Nat. Publ. Gr. 13, 370–383 (2016).

11. Piraner, D. I. et al. Going Deeper: Biomolecular Tools for Acoustic and Magnetic Imaging and Control of Cellular Function. (2017). doi:10.1021/acs.biochem.7b00443

12. Rev, A. et al. Biomolecular Ultrasound and Sonogenetics. (2018).

13. Shapiro, M. G. et al. Biogenic gas nanostructures as ultrasonic molecular reporters. Nat. Nanotechnol. 9, 311–6 (2014).

14. Lu, G. J. et al. imaging of gas-filled protein nanostructures. Nat. Mater. 17, (2018).

15. Bourdeau, R. W. et al. Acoustic reporter genes for noninvasive imaging of microorganisms in mammalian hosts. Nature 553, 86–90 (2018).

16. Shapiro, M. G. et al. Genetically encoded reporters for hyperpolarized xenon magnetic resonance imaging. Nat. Chem. (2014). doi:10.1038/nchem.1934

17. Pfeifer, F. Distribution, formation and regulation of gas vesicles. Nat. Rev. Microbiol. 10, 705–715 (2012).

18. Walsby, A. E. Gas Vesicles. Annu. Rev. Plant Physiol. 26, 427–439 (1975).

19. Lakshmanan, A. et al. Molecular Engineering of Acoustic Protein Nanostructures. ACS Nano 10, 7314–7322 (2016).

20. Kunth, M., Lu, G. J., Witte, C., Shapiro, M. G. & Schro, L. Protein Nanostructures Produce Self-Adjusting Hyperpolarized Magnetic Resonance Imaging Contrast through Physical Gas Partitioning. (2018). doi:10.1021/acsnano.8b04222

21. Maresca, D., Sawyer, D. P., Renaud, G., Lee-gosselin, A. & Shapiro, M. G. Nonlinear X-Wave Ultrasound Imaging of Acoustic Biomolecules. Phys. Rev. X 8, 41002 (2018).

22. Maresca, D. et al. Nonlinear ultrasound imaging of nanoscale acoustic biomolecules Nonlinear ultrasound imaging of nanoscale acoustic biomolecules. 073704, (2017).

23. Ferrara, K., Pollard, R. & Borden, M. Ultrasound Microbubble Contrast Agents : Fundamentals and Application to Gene and Drug Delivery. (2007). doi:10.1146/annurev.bioeng.8.061505.095852

24. Ashokkumar, M., Lee, J., Kentish, S. & Grieser, F. Bubbles in an acoustic field : An overview. 14, 470–475 (2007).

25. Kwan, J. J. et al. Ultrasound-Propelled Nanocups for Drug Delivery. Small 11, 5305–5314 (2015).

26. Coussios, C. C. & Roy, R. A. Applications of Acoustics and Cavitation to Noninvasive Therapy and Drug Delivery. (2008). doi:10.1146/annurev.fluid.40.111406.102116

27. Holland, C. K. & Apfel, R. E. An Improved Theory for the Prediction of Microcavitation Thresholds. 204–208 (1989).

28. Church, C. C. & Church, C. C. Frequency, pulse length, and the mechanical index. 162, 1–8 (2007).

29. Cherin, E. et al. Acoustic behavior of Halobacterium salinarum gas vesicles in the high-frequency range: experiments and modeling. Ultrasound Med. Biol. 43, 1016–1030 (2017).

30. Walsby, A. E. The pressure relationships of gas vacuoles. Proc. R. Soc. London. Ser. B. Biol. Sci. 178, 301–326 (1971).

31. Lakshmanan, A. et al. Preparation of biogenic gas vesicle nanostructures for use as contrast agents for ultrasound and MRI. Nat. Protoc. 12, 2050–2080 (2017).

32. Häcker, G., Redecke, V. & Häcker, H. Activation of the immune system by bacterial CpG-DNA. Immunology 105, 245–251 (2002).

33. Dang, L. H., Bettegowda, C., Huso, D. L., Kinzler, K. W. & Vogelstein, B. Combination bacteriolytic therapy for the treatment of experimental tumors. Proc. Natl. Acad. Sci. U. S. A. 98, 15155–60 (2001).

34. Ryan, R. M. et al. Bacterial delivery of a novel cytolysin to hypoxic areas of solid tumors. Gene Ther. 16, 329–339 (2009).

35. Forbes, N. S., Munn, L. L., Fukumura, D. & Jain, R. K. Sparse initial entrapment of systemically injected Salmonella typhimurium leads to heterogeneous accumulation within tumors. Cancer Res. 63, 5188–5193 (2003).

36. Cobbold, R. S. Foundations of biomedical ultrasound. (Oxford University Press, 2006).

37. Zhang, H. F., Maslov, K., Sivaramakrishnan, M., Stoica, G. & Wang, L. V. Imaging of hemoglobin oxygen saturation variations in single vessels in vivo using photoacoustic microscopy. Appl. Phys. Lett. 90, 5–7 (2007).

38. Beard, P. Biomedical photoacoustic imaging. Interface Focus 1, 602–631 (2011).

39. Jang, M. J. & Nam, Y. NeuroCa: integrated framework for systematic analysis of spatiotemporal neuronal activity patterns from large-scale optical recording data. Neurophotonics 2, 035003 (2015).

