## Supplementary Tables, Figures, and Movie Captions for "Acoustically Detonated Biomolecules for Genetically Encodable Inertial Cavitation"

### Supplementary Tables, Figures and Movies

| Detected Harmonic # | Frequency ($f_0$ = 0.67 MHz) | Frequency ($f_0$ = 3 MHz) |
|---|---|---|
| 1 | 8.04 | 8.96 |
| 2 | 8.71 | 11.93 |
| 3 | 9.38 | 14.95 |
| 4 | 10.05 | 17.93 |
| 5 | 10.72 | 20.86 |
| 6 | 11.40 | 23.83 |
| 7 | 12.06 | |
| 8 | 12.73 | |
| 9 | 13.40 | |
| 10 | 14.08 | |
| 11 | 14.74 | |
| 12 | 15.42 | |
| 13 | 16.08 | |
| 14 | 16.75 | |
| 15 | 17.43 | |
| 16 | 18.09 | |
| 17 | 18.76 | |
| 18 | 19.43 | |
| 19 | 20.11 | |
| 20 | 20.77 | |
| 21 | 21.44 | |
| 22 | 22.11 | |
| 23 | 22.78 | |
| 24 | 23.45 | |
| 25 | 24.12 | |
| 26 | 24.79 | |
| 27 | 25.46 | |
| 28 | 26.13 | |
| 29 | 26.81 | |

**Supplementary Table 1 | Frequencies of peak harmonic signals detected by PCD.** Harmonics of the central frequency $f_0$ = 0.67MHz are presented in the middle, and harmonics of the central frequency $f_0$ = 3MHz are presented on the right.

 All rights reserved. 

| Plasmid | GV cassette | Payload |
| --- | --- | --- |
| pABZ_01 | ARG1 (A2C) | NanoLuc |
| pABZ_02 | --- | NanoLuc |

**Supplementary Table 2 | Genetic constructs used in this study.** All plasmids were constructed using the pTD103 backbone with a NanoLuc protein payload, with or without a GV-producing gene cassette. Sources of genetic elements: pTD103 plasmids: J. Hasty, UCSD; ARG1: Addgene #106473; NanoLuc: Addgene #87696

 All rights reserved. 

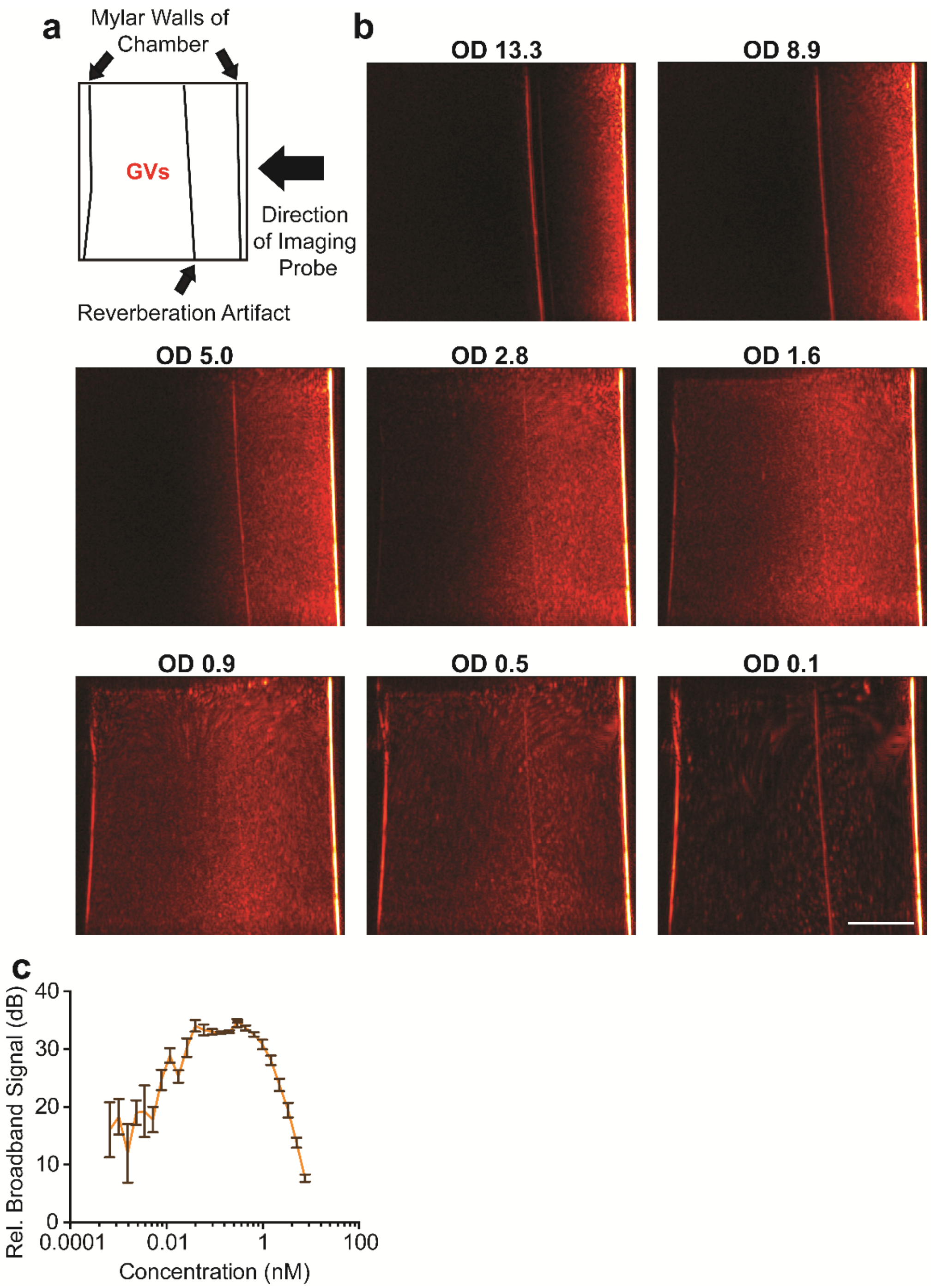


**Supplementary Figure 1 | GVs attenuate ultrasound at high concentrations. a**, Illustration of the sample chamber and setup as seen in the images. **b**, B-mode images of purified Ana GVs in different concentrations showing acoustic shadowing in high concentrations. Scale bar represents 3 mm. **c**, The effect of high GV concentrations on average broadband measurements. GVs were insonated with a single 30-cycle pulse with PNP = 1.0 MPa (n=5).

 All rights reserved. 

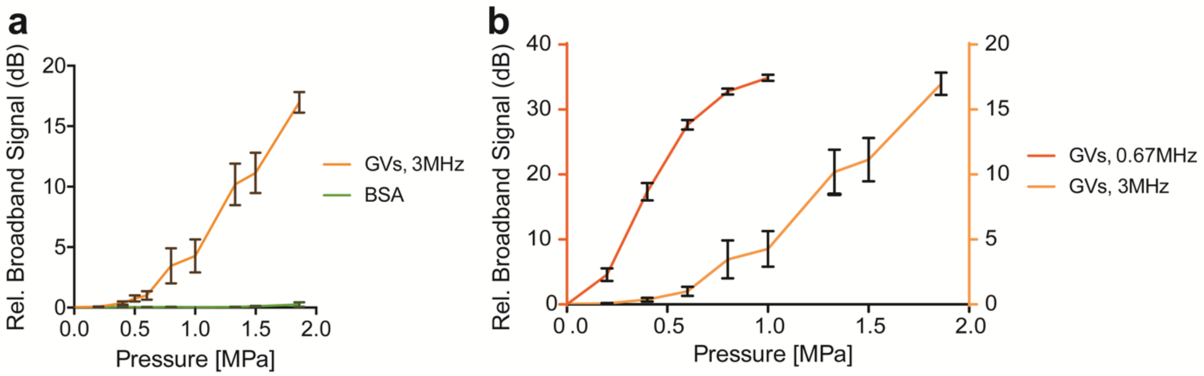


**Supplementary Figure 2 | GV-seeded cavitation at 3 MHz requires higher pressure levels. a**, Broadband signals recorded from GVs (0.3 nM) and BSA (matched in mg/mL to GVs concentration) insonated at 3MHz. Broadband signal increased with pressure and was significantly higher for GV samples for PNP >= 0.5 MPa ($p < 0.05$ for PNP < 1.33 MPa, and $p < 0.001$ at higher pressure levels, $n = 8$). **b**, Comparison between broadband signals from GVs insonated with 0.67 MHz and 3 MHz pulses.

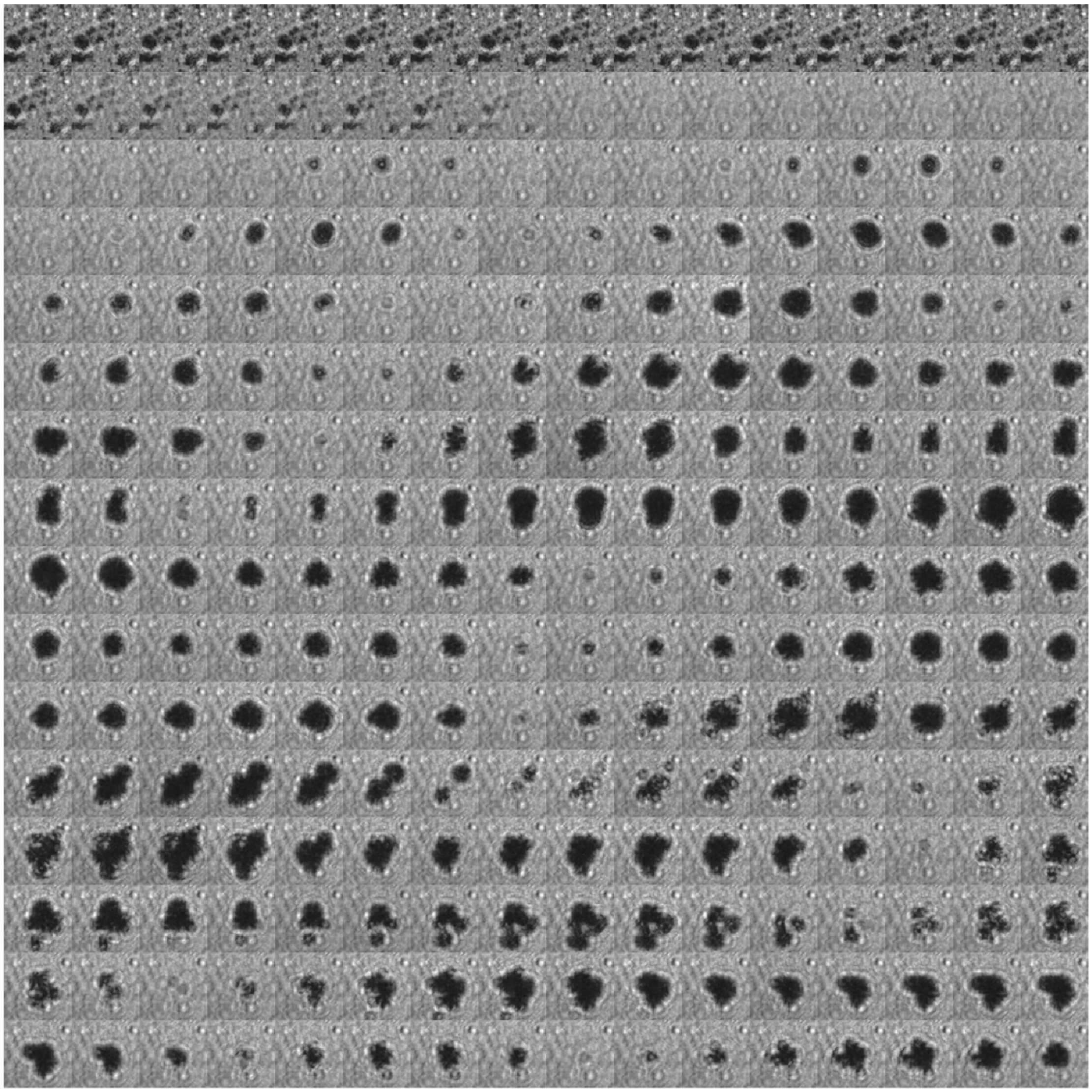

**Supplementary Figure 3 | High frame rate optical imaging of GV collapse and bubble cavitation.** High-speed camera frames (left to right then top to bottom) of GV collapse and cavitation (200 ns between each frame, 31x31 µm field of view), focusing on a single bubble. Initial black spots are GVs, which disappear due to collapse, and a cavitating bubble then appears in this region.

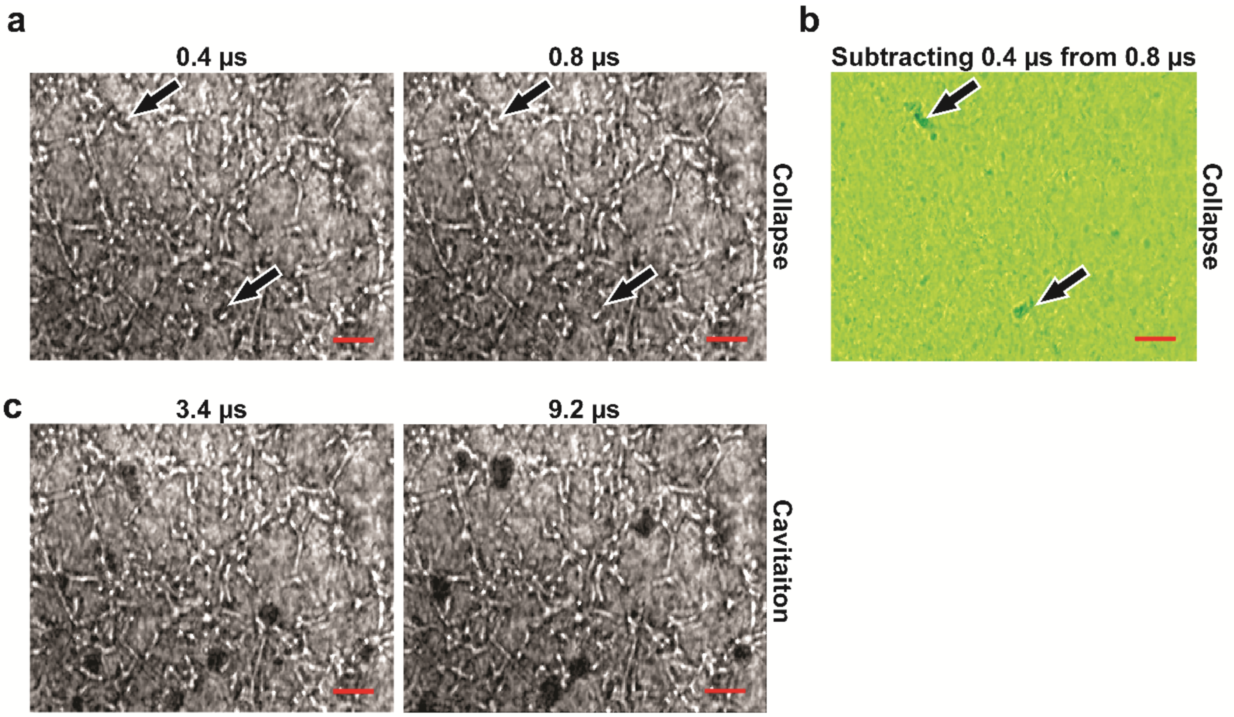


**Supplementary Figure 4 | High frame rate recording of GVs attached to tumor cells.** GVs attached to U87 cells (0.4 µs) are collapsed by the ultrasound wave (0.8 µs, and differential image). Only after the collapse of the GVs are cavitation events seen (3.4 µs and 9.2 µs).

**Supplementary Movie 1 | Representative high frame rate movie of GV attached to a Mylar plate.** A series of 256 images showing cavitation nucleated by GVs attached to a Mylar plate were collected over 51.2 µsec at 5 million frames per second (fps) The movie is displayed at 5 fps, one million times slower than the real time.

**Supplementary Movie 2 | Representative high frame rate movie of GVs attached to tumor cells.** A series of 256 images showing cavitation nucleated by GVs attached to U87 tumor cells were collected over 51.2 µsec at 5 million frames per second (fps) The movie is displayed at 5 fps, one million times slower than the real time.


bioRxiv preprint. April 26, 2019. All rights reserved.
